# Updates on enzymatic and structural properties of human glutamine: fructose-6-phosphate amidotransferase 2 (hGFAT2)

**DOI:** 10.1101/2020.07.13.201285

**Authors:** Isadora A. Oliveira, Diego Allonso, Tácio V. A. Fernandes, Daniela M. S. Lucena, Gustavo T. Ventura, Wagner B. Dias, Ronaldo S. Mohana-Borges, Pedro G. Pascutti, Adriane R. Todeschini

**Affiliations:** Laboratório de Glicobiologia Estrutural e Funcional, Instituto de Biofísica Carlos Chagas Filho (IBCCF), Universidade Federal do Rio de Janeiro (UFRJ); Departamento de Biotecnologia Farmacêutica, Faculdade de Farmácia, UFRJ; Laboratório de Modelagem e Dinâmica Molecular, IBCCF, UFRJ; Laboratório de Macromoléculas, Diretoria de Metrologia Aplicada às Ciências da Vida, Instituto Nacional de Metrologia, Qualidade e Tecnologia (INMETRO), Duque de Caxias, RJ 25250-020, Brazil; Laboratório de Genômica Estrutural, IBCCF, UFRJ, Rio de Janeiro, RJ 21941-902, Brazil

**Keywords:** Glutamine:fructose-6-phosphate amidotransferase (GFAT), glucosamine-6-phosphate synthase, hexosamine biosynthetic pathway (HBP), carbohydrate metabolism, enzyme kinetics, protein structure, molecular modelling, molecular dynamics

## Abstract

Glycoconjugates play a central role in several cellular processes and alteration in their composition is associated to human pathologies. The hexosamine biosynthetic pathway is a route through which cells obtain substrates for cellular glycosylation, and is controlled by the glutamine: fructose-6-phosphate amidotransferase (GFAT). Human isoform 2 GFAT (hGFAT2) has been implicated in diabetes and cancer, however, there is no information about structural and enzymatic properties of this enzyme. Here, we report a successful expression and purification of a catalytically active recombinant hGFAT2 (rhGFAT2) in *E. coli* cells fused or not to a HisTag at the C-terminal end. Our enzyme kinetics data suggest that hGFAT2 does not follow the ordered bi-bi mechanism, and performs the glucosamine-6-phosphate synthesis much slowly than previously reported for other GFATs. In addition, hGFAT2 is able to isomerase fructose-6-phosphate into glucose-6-phosphate even in presence of equimolar amounts of glutamine, in an unproductive glutamine hydrolysis. Structural analysis of the generated three-dimensional model rhGFAT2, corroborated by circular dichroism data, indicated the presence of a partially structured loop in glutaminase domain, whose sequence is present in eukaryotic enzymes but absent in the *E. coli* homolog. Molecular dynamics simulations show such loop as the most flexible portion of the protein, which interacts with the protein mainly through the interdomain region, and plays a key role on conformational states of hGFAT2. Altogether, our study provides the first comprehensive set of data on the structure, kinetics and mechanics of hGFAT2, which will certainly contribute for further studies focusing on drug development targeting hGFAT2.

## Introduction

Glycoconjugates are particularly diverse in structure and composition and play a central role in several cellular processes such as cell growth, cell-cell and cell-matrix adhesion, cell differentiation among others. Severe alterations in the composition of glycoconjugates are usually associated to human diseases (1,2). The primary substrates for intra- and extracellular glycosylation are obtained through the hexosamine biosynthetic pathway (HBP), which is controlled by the rate-limiting enzyme glutamine:fructose-6-phosphate amidotransferase (GFAT) (3).

The enzyme GFAT belongs to the amidotransferase family, class II, characterized by an N-terminal cysteine as the nucleophilic catalyst (4). All cellular organisms including prokaryotes and eukaryotes express this class of enzymes, highlighting their relevance to normal cell functioning. Indeed, deletion of the GFAT gene in *Escherichia coli* and *Saccharomyces pombe* led to cell death (5). In mammals, GFAT was characterized in 1960 in rat liver homogenates, when Ghosh et al. (6) described its specificity for fructose-6-phosphate (Fru-6P) and not glucose-6-phosphate (Glc-6P) for glucosamine-6-phosphate (GlcN-6P) generation. In humans, three different isoforms of GFAT were reported, named hGFAT1, hGFAT1Alt (or GFAT1-L) and hGFAT2, encoded by the *gfpt1* and *gfpt2* genes, respectively. hGFAT1 expression is ubiquitous and it is highly expressed in placenta, pancreas and testis (7). hGFAT1Alt represents an expanded isoform of hGFAT1 resulting from alternative splicing of the *gfpt1* gene, and its expression is restricted to striated muscle (8,9). In turn, hGFAT2 is the product of a distinct gene, *gfpt2*, and shares 79% identity with hGFAT1 (7). hGFAT2 presents a more restricted expression pattern than hGFAT1, being the major isoform in several central nervous system tissues and observed in a smaller proportion in the heart, placenta, testis and ovary (7).

Interest in hGFAT has increased in the past few years as this protein has been implicated in human pathologies. The hGFATs play a direct role in type 2 diabetes and their overexpression contributes to insulin resistance and higher *O*-GlcNAc levels (10–12). In fact, previous work has identified hGFATs as potential targets for the development of anti-diabetes drugs (12,13). In addition to diabetes, hGFAT has been assigned a prominent role in the close relationship between HBP and cancer. hGFAT1 isoform has been observed to be up-regulated in breast (14), prostate (15) and hepatic (16) cancers. On the other hand, it has been observed that hGFAT2 levels increased considerably in pancreatic adenocarcinoma (17) and colorectal cancer (18).

Despite its importance in cellular metabolism, there is few works uncovering biochemical and kinetics properties of mammalian GFATs. The structure of GFAT is characterized by having two domains, glutaminase (GLN) and isomerase (ISOM), responsible for its enzymatic activity. The complete reaction mechanism of GFAT was proposed from studies with GlcN-6P synthase (GlmS), the bacterial homolog of hGFAT, being characterized as bi-bi-ordered in which the entry of Fru-6P induces conformational changes that favor glutamine (Gln) binding (19). Concerning the hGFATs, most studies to date have focused on unraveling the mechanisms and structure of isoform 1 (20–22). This isoform naturally occurs as a homotetramer, which is its active oligomeric state (21,23). In contrast to *E. coli* GlmS, there were few crystal structures of the hGFAT1 isomerase (ISOM) domain, and only very recently the full structure of this isoform was reported (22). Conversely, there is only one report focusing on the expression, purification and kinetics of the recombinant variation of the murine GFAT2 (mGFAT2) (24).

Therefore, studies concerning the characterization of physical-chemical properties of hGFAT2 are certainly needed. In the present work, we successfully expressed and purified the hGFAT2 in *E. coli* cells with and without a HisTag at C-terminal end, and demonstrated that hGFAT2 forms a tetramer in solution with full catalytic activity in both constructs. However, hGFAT2 exhibits a lower catalytic efficiency than those reported from other homologs or hGFAT1, but similar *K*_*M*_ values. Using a series of enzyme assays, we detected a decoupling between the activities of both GFAT2 domains regardless the HisTag, generating ammonia leakage even in presence of Gln, and Fru-6P isomerization in Glc-6P. Despite the high identity between hGFAT1 and hGFAT2, the last is, in contrast, poorly inhibited by UDP-GlcNAc. Molecular dynamics simulations of the modeled hGFAT2 structure suggests the UDP-GlcNAc binding pocket may be partially blocked by interactions between the interdomain region and a highly flexible loop, whose structure was not elucidated in hGFAT1.

## Results

### Recombinant human GFAT2 (rhGFAT2) forms tetramers in solution

Our current knowledge on hGFAT rests on few studies that focused on hGFAT1 (20–22). However, recent evidences suggest an important association between hGFAT2 and metabolic pathologies (18,25). In this context, a comprehensive characterization of hGFAT2 has become more pressing.

As previous literature diverges on the impact of fusion tags in both GlmS and GFAT activity (21), we expressed the recombinant hGFAT2 (rhGFAT2) protein in *E. coli* cells with and without a 6xHisTag at its C-terminal end. The best expression condition for both constructs was achieved after 0.5 mM IPTG induction for 6 h at 25 °C under agitation. Although the majority of rhGFAT2 was expressed as inclusion bodies, a small fraction remained soluble (Suppl. Fig 1A. and B). To avoid improper refolding, we purified rhGFAT2 from the soluble fraction in a Ni^+2^NTA column and we obtained highly pure (96.6% purity) HisTag-containing rhGFAT2 (rhGFAT2-his) protein after a single step of affinity chromatography (Suppl. Fig. 1A, Table 1). Surprisingly, rhGFAT2 without HisTag (rhGFAT2 w/o tag) also bound to Ni^+2^NTA column, comprising therefore the first step for its purification, which reached high purity level (96.3%) after an additional step of an anion exchange chromatography in a Q-sepharose column (Suppl. Fig. 1B, Table 1). Despite the high purity of both samples, final yield of purified rhGFAT2 w/o tag was 10 times lower than that of rhGFAT2-his (Table 1). The absence of the HisTag was further confirmed by Western blot analysis using an anti-HisTag monoclonal antibody (Suppl. Fig. 1C).

**Table 1.**
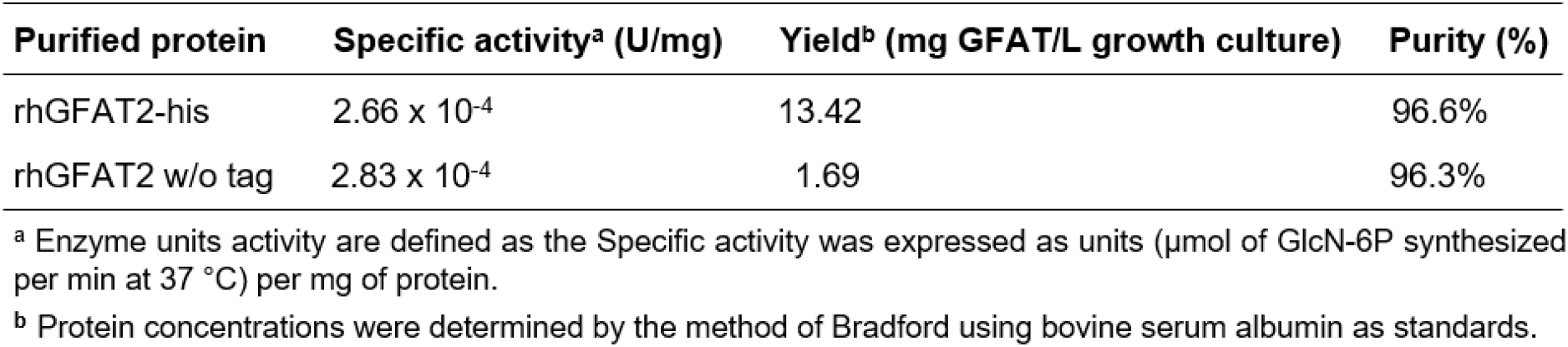
Purification of rhGFAT2s in *E. coli*.

| Purified protein | Specific activity<sup>a</sup> (U/mg) | Yield<sup>b</sup> (mg GFAT/L growth culture) | Purity (%) |
| --- | --- | --- | --- |
| rhGFAT2-his | 2.66 x 10<sup>-4</sup> | 13.42 | 96.6% |
| rhGFAT2 w/o tag | 2.83 x 10<sup>-4</sup> | 1.69 | 96.3% |

<sup>a</sup> Enzyme units activity are defined as the Specific activity was expressed as units (μmol of GlcN-6P synthesized per min at 37 °C) per mg of protein.
<sup>b</sup> Protein concentrations were determined by the method of Bradford using bovine serum albumin as standards.

To evaluate the enzymatic activity of the purified enzymes, we performed an enzymatic assay to detect the GlcN-6P formation. As shown in Table 1, both rhGFAT2-his and rhGFAT2 w/o tag exhibited similar specific activity (Table 1), suggesting that HisTag at the C-terminal end has no effect on enzyme function. Based on these results and overall yield of purified enzymes, we decided to set rhGFAT2-his as the target for further biochemical characterization studies.

Structural data regarding hGFAT1 indicates that this protein is found as a homotetramer in solution (21,23). To assess whether rhGFAT2 also forms oligomers, we performed a cross-link assay using ethylene glycol bis(succinimidyl succinate) (EGS) as cross-linking agent and different amounts of rhGFAT2. We observed the presence of tetramers in all conditions analyzed, however we only achieved complete tetramerization when using 10 μg of rhGFAT2 (Fig. 1A). To confirm this finding, we also performed a size exclusion chromatography using the Superdex 200 column. The fractions containing rhGFAT2 were eluted with the retention volume of 95 mL (Fig. 1B and C), indicating a molecular weight of approximately 300 kDa, which is consistent with the expected molecular weight of the rhGFAT2 tetramer. It is worth noting that an additional peak observed at approximately 74 mL corresponds to higher oligomers, possibly octamers (Fig. 1B and C). Altogether, our results demonstrate that rhGFAT2 can be successfully expressed in *E. coli* cells and the purified protein forms tetramers in solution.

**Fig. 1.**
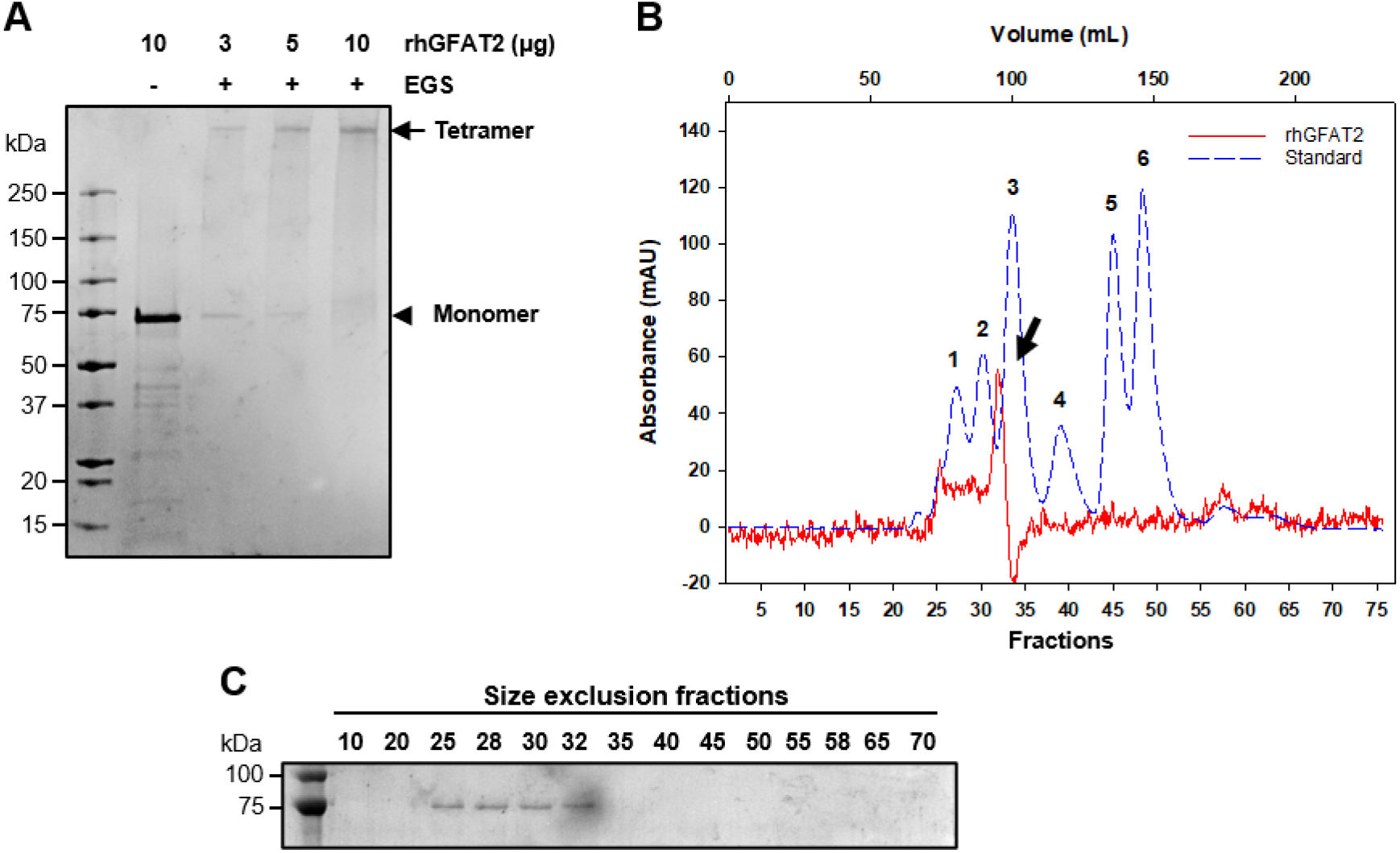
Evaluation of the rhGFAT2 oligomeric state. (A) Cross-linking assay in which 3, 5, or 10 μg of hGFAT2-his were incubated in the presence of 1 mM EGS. The control was performed by incubation of 10 μg of rhGFAT2-his in the absence of EGS. Arrow and arrowhead represent the tetramer and monomer, respectively. (B) Size exclusion chromatogram of rhGFAT2-his (solid red line) in Superdex 200 16/200 column. The arrow represents the major peak of the enzyme. Standard proteins (dashed blue line) were subjected to the same condition as rhGFAT2 and are described as follows: 1-Thyroglobulin (669 kDa), 2-Apoferritin (443 kDa), 3-β-amylase (200 kDa), 4-BSA (66 kDa), 5-Carbon anhydrase (29 kDa) and 6-Cytochrome c oxidase (12.4 kDa). The collected fractions were subjected to SDS-PAGE followed by Coomassie blue staining (C).

### Enzyme kinetics of rhGFAT2

In order to have a detailed perspective on hGFAT2 kinetics, we have measured its GlcN-6P synthetic activity (Suppl. Fig. 2A), using a modified Elson-Morgan reaction (26,27), and the ability of each domain to hydrolyze Gln or isomerase Fru-6P through coupled assays. All kinetic parameters are summarized in Table 2. Since the GlcN-6P synthetic activity of hGFAT is suggested to follow a bisubstrate ordered mechanism based on kinetic studies of *E. coli* GlmS (19,28), we initially used that mechanistic model equation to fit our rate *versus* substrate curves (Suppl. Fig. 2A). However, we obtained an inconsistent (negative) value for ^*Fru-6P*^*K*_*M*_. Hence, we used Michaelis-Menten model to obtain apparent values of *K*_*M*_ and *k*_*cat*_. The ^app^*K*_*M*_ value for Fru-6P is higher than for Gln (0.957 and 0.763 mM, respectively) (Table 2). Noteworthy, hGFAT2-his exhibited a low ^app^*k*_*cat*_ value for GlcN-6P synthesis (Table 2). Since the presence of HisTag did not affected the overall synthetic activity (Table 1), we checked if the N-terminal methionine (Met1) was removed during heterologous expression, which could hamper the Cys2 to perform its role as catalyst and could explain the reduced activity of hGFAT2-his.

**Table 2.** Kinetic parameters of relations catalysed by rhGFAT2-his.

| Type of activity | Substrate(s) | $K_m$ (Gln) (mM) | $K_m$ (Fru-6P) (mM) | $k_{cat}$ (min<sup>-1</sup>) |
| --- | --- | --- | --- | --- |
| Aminohydrolysing activity | w/o Fru-6P | 0.820±0.335 | – | 0.021±0.003 |
|  | w/ Fru-6P | 1.814±1.158 | – | 0.079±0.016 |
| Isomerase activity | w/o Gln | – | 0.711±0.170 | 0.322±0.022 |
| GlcN-6P synthetic activity | ▲ Fru-6P | – | 0.957±0.502* | 0.032±0.007* |
|  | ▲ Gln | 0.763±0.332* | – | 0.040 ±0.008* |

\* The kinetic parameters for synthase activity were generated through Michaelis-Menten fitting, therefore must be considered as apparent values.

However, the peptide fingerprint suggests the Met1 was properly removed, as observed in the coverage of the peptides detected (Suppl. Fig. 3A), and confirmed by the fragmentation pattern of the N-terminal peptide 2-CGIFAYMNYRVPR-14 (Suppl. Fig. 3B).

Using a coupled assay, we monitored the release of glutamic acid from Gln hydrolysis, by which we observed that hGFAT2-his is able to execute such activity even in absence of Fru-6P, but the presence of this phosphorylated monosaccharide increases 4 times the *k*_*cat*_ of aminohydrolysis reaction of (Table 2). The kinetic curves for the Gln hydrolysis show the increase in the rate bursted by Fru-6P (Suppl. Fig 2B), with a direct impact on the catalytic efficiency (Table 2). Whereas *k*_*cat*_ of hGFAT2-his aminohydrolysing activity is in the same order of magnitude as of synthetic activity (around 0.03 min^−1^), the ISOM activity exhibits a 10-times higher *k*_*cat*_ (0.322 min^−1^), and a *K*_*M*_ of 0.711 mM for the sugar (Table 2). In contrast to aminohydrolysis, the analysis of the ISOM activity curves in the presence of increasing concentrations of Gln indicates that hGFAT2-his performs the isomerization of Fru-6P to Glc-6P even in higher concentrations of this amino acid (Suppl. Fig 2C), suggesting that part of the ammonia released from Gln hydrolysis is not used for GlcN-6P synthesis.

In order to confirm the unproductive hydrolysis of Gln, we used nuclear magnetic resonance (NMR) to detect directly the substrate consumption and the product formation from hGFAT2-his activity (Fig. 2A). Equimolar amounts (3 mM) of Gln and Fru-6P was incubated in the presence or absence of hGFAT2-his and the 1D ^1^H spectra were acquired (Supp. Fig. 4A and 4B). As expected, we observed the consumption of both Gln (reduced peaks at 2.15 and 2.48 ppm, corresponding to Hβ and Hγ, Fig. 2B) and Fru-6P (reduced peaks at 4.25 and 4.17 ppm, corresponding to H1 and H3, Fig. 2C) concomitantly with the generation of Glu (increased peaks at 2.07 and 2.36 ppm, corresponding to Hβ and Hγ, Fig. 2B) and αGlcN-6P (increased peaks at 5.42 and 4.062 ppm, corresponding to H1 and H6, Fig. 2C) catalyzed by rhGFAT2-his. Noteworthy we noticed that peaks from both Glc-6P anomers increased during the reaction time course (5.23, 3.28, 3.52 and 4.00 ppm corresponding to αH1, βH2, βH3 and βH4, respectively, Fig. 2C). The H1 from β-sugars were not detected, probably due to distortion of the spectra by the water suppression at 4.70 ppm. However, TOCSY spectrum at t = 84 h exhibits the correlation signals among H1, H2 and H3 from βGlc-6P (4.65, 3.28 and 3.52 ppm, respectively, Suppl. Fig. 4C). The TOCSY spectra also exhibits the correlation signals among Hα, Hβ and Hγ from both Gln and Glu (Suppl. Fig. 4C). Although close to the noise, the correlation signals between H1 and H3 from αGlcN-6P (5.42 and 3.93 ppm, respectively), and among H1, H3 and H5 from αGlc-6P (5.23, 3.75 and 3.92 ppm, respectively) were also observed (data not shown). The αH1 signal from Glc-6P is present in a proportion of 1:2.5 relative to αH1 of GlcN-6P measured in 1D ^1^H spectra, showing that rhGFAT2 partially acts as an isomerase even at equimolar concentrations of both substrates, corroborating the ISOM kinetics data. In addition, we did not observe spontaneous isomerization from Fru-6P to Glc-6P, spontaneous hydrolysis of Gln or GlcN-6P formation in the absence of the enzyme (Suppl. Fig. 4B). The oxidation of DTT (29) was the sole alteration in 1D ^1^H spectrum observed in the absence of the enzyme (Suppl. Fig. 4B).

**Fig. 2.**
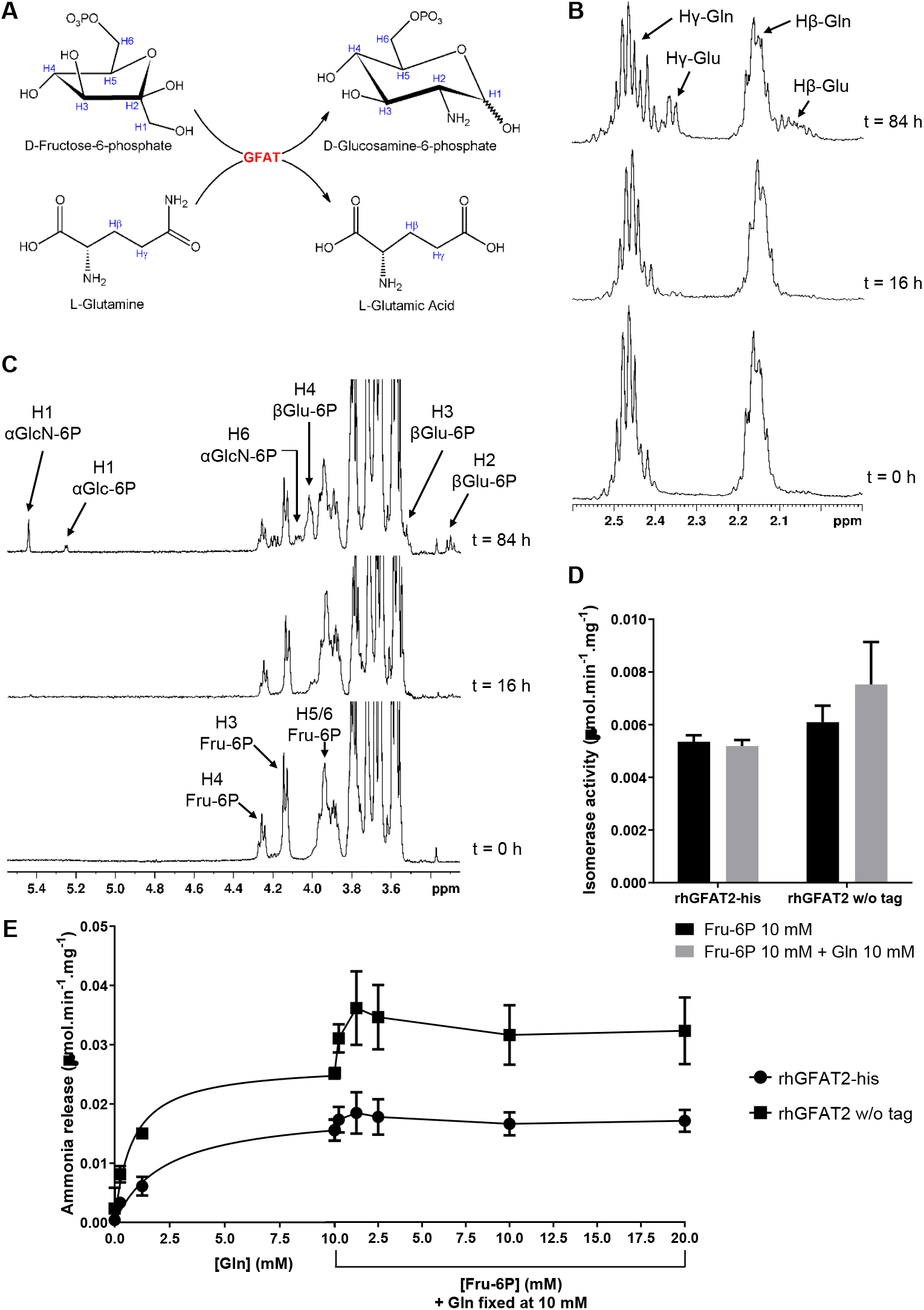
Exploring PGI-like activity of rhGFAT2. (A) Reaction scheme of complete GFAT reaction. (B and C) Time course of rhGFAT2-his reaction in presence of Gln and Fru-6P (both at 3 mM) in 50 mM deuterated phosphate buffer pH 7.4 with 1 mM DTT at 25 °C, monitored by ^1^H NMR spectroscopy. The times at which spectral data were acquired refer to the addition of rhGFAT2-his (100 μg) as t = 0 h. (B) Region of ^1^H NMR spectra detailing Gln and Glu peaks. (C) Region of ^1^H NMR spectra detailing Fru-6P, Glc-6P and GlcN-6P peaks. (D) Isomerization of Fru-6P in Glc-6P catalyzed by rhGFAT2-his or rhGFAT2 without (w/o) HisTag assessed either in absence (black bars) or in presence (gray bars) of Gln. (E) Ammonia release from Gln, catalyzed by rhGFAT2-his (black circles) and rhGFAT2 without (w/o) HisTag (black squares). The enzyme was incubated with increasing amounts of Gln until 10 mM, and with fixed Gln at 10 mM and variable concentrations of Fru-6P as indicated.

In order to evaluate whether the innocuous effect of Gln on ISOM activity (Suppl. Fig. 2C) was due the presence of the C-terminal HisTag, we performed such assay with rhGFAT2 without tag. As observed for rhGFAT2-his, the addition of Gln in reaction medium did not only reduce the ISOM activity of rhGFAT2 w/o tag (Fig. 2D), but even enhances it suggesting that part of the ammonium released from Gln hydrolysis in GLN domain does not reach the ISOM domain.

To access the ammonia release from glutamine hydrolysis catalyzed by GFAT, we used a coupled assay with glutamic acid dehydrogenase in the presence of α-ketoglutaric acid and NADH. The reduction in NADH absorbance was correlated to ammonia release using a standard curve of NH_4_Cl. As we can see in Fig. 2E, the ammonia release increases with increasing concentrations of Gln, as expected, for both rhGFAT2 with and without HisTag. Fitting the curves with Michaelis-Menten equation, we observed that both enzymes reached a plateau at Gln saturating concentration (10 mM), but the ammonia release from rhGFAT2 w/o tag is twice the values obtained from the HisTaggeg enzyme. Furthermore, the addition of Fru-6P, even at high concentrations, did not abolish the ammonia released to the medium for both the enzymes (Fig. 2E), but actually enhancing the ammonia release, mainly for rhGFAT2 w/o tag. These data indicate that a great amount of the ammonia hydrolyzed from Gln is lost to the medium instead of reaching the ISOM domain for generating GlcN-6P.

### rhGFAT2 inhibition by UDP-GlcNAc

UDP-GlcNAc, the final product of HBP, has been described as a potent inhibitor of glutaminase activity of hGFAT1 (20,22). To examine whether UDP-GlcNAc is able to inhibit hGFAT2 as well, we assayed rhGFAT2’s glutaminase activity in the presence of different concentrations of the activated monosaccharide. By plotting the results in a Dixon plot (Suppl. Fig. 5), we observed that UDP-GlcNAc is able to inhibit only 10% of rhGFAT2 activity, behaving as a partial inhibitor. This result shows that UDP-GlcNAc is a weak inhibitor of this GFAT isoform.

### Unstructured loop as a key for interdomain (miss)communication

In an effort to understand the differences among the kinetic data reported for other GFATs and our results, we compared the sequences of GlmS (GFAT from *E. coli*), GFA (GFAT from *C. albicans*), hGFAT1 and hGFAT2. The alignment between GlmS and the hGFATs showed that the human variants exhibit an additional 46 residues sequence, from Lys211 to Val256 (hGFAT2 numbering) (Fig. 3A). This internal sequence is also present in GFA and is longer than those from hGFATs (Fig. 3A). Besides, these additional sequences are the most distinctive portion among hGFATs and GFA, and even between hGFAT1 and hGFAT2 (Fig. 3A): the identity of these short sequences between the human isoforms is only 49%, in contrast to 79% when we compare the full sequences.

**Fig. 3.**
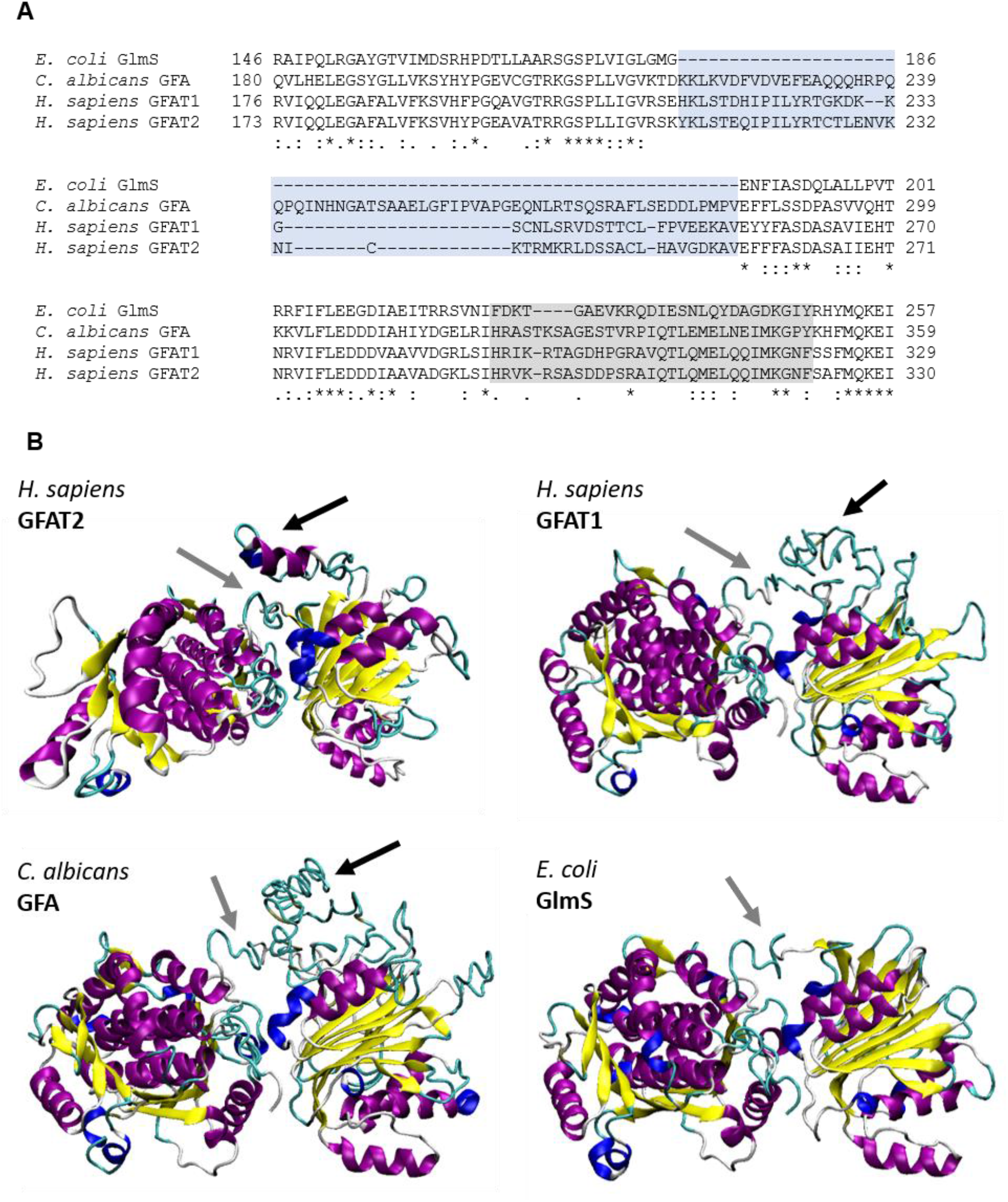
Structure insights on human, fungal and bacterial GFATs. (A) Alignment of the internal loop and interdomain connection sequences (highlighted in blue and gray, respectively) from GlmS (*E. coli*), GFA (*C. albicans*), and GFAT1 and GFAT2 (*H. sapiens)* performed with Clustal Omega server. Dashes indicate gaps, asteriscs indicate identical residues, and dots indicate residues with similar physical-chemical properties. (B) Three-dimensional models obtained for hGFAT2, hGFAT1, and GFA from *C. albicans* from threading using I-TASSER server. The structure of GlmS was retrieved from PDB under ID 4AMV. The proteins are represented in cartoon and colored according their secondary structure (α-helix in purple, β-sheet in yellow, 3-10 helix in blue, turns in cyan, and coil in white). The black arrows indicate the loop regions; the gray arrows indicate the interdomain region.

To better analyze the impact of these internal sequences on GFATs’ structures, we generated tridimensional model for hGFAT2 using threading methods by subjecting the hGFAT2 amino acid sequence to I-TASSER and LOMETS servers. We opted, therefore, for the I-TASSER final model because it presented the best loop conformation. In this model, part of the loop (black arrow) folded into an α-helix structure due to the interaction with intra-chain residues and the stabilization occurred by interaction with the interdomain connective portion (gray arrow, Fig. 3B). Although homology modelling could have used for this task, especially given the recently available full structure of hGFAT1 (22), there is a lack of structural information regarding exactly the sequence Lys211-Val256 throughout the experimental structures available. In addition, the model reliability is corroborated by the similar secondary structure content observed experimentally by circular dichroism for rhGFAT2-his (Suppl. Fig. 6, Table 3).

**Table 3.** Secondary structure composition of rhGFAT2, based on experimental and theoretical data.

| Method | α-Helix | β-Sheet | Random |
| --- | --- | --- | --- |
| Circular Dichroism<sup>a</sup> | 33% | 18% | 49% |
| Molecular Dynamics<sup>b</sup> | 36% | 19% | 45% |

<sup>a</sup> The secondary structure content of rhGFAT2-his was estimated from circular dichroism data, using different algorithms available on the Dichroweb server.
<sup>b</sup> The average secondary structure of GFAT2 from molecular dynamics simulation time was shown.

By using the same threading approach for hGFAT1 and GFA from *C. albicans*, we obtained the models for the complete structure of both. As shown in Fig. 3B, the additional sequence of hGFAT1 also forms an unstructured loop, similar to that from hGFAT2, close to the interdomain region. Whereas bigger and even less structured, the loop from GFA is also next to interdomain connective portion, contrasting to GlmS structure (PDB ID: 4AMV), in which such a loop is absent (Fig. 3B).

These results prompted us to investigate a possible function for the loop. Thus, we performed 3-replica molecular dynamics (MD) simulations of 500 ns each using the AMBER package. In fact, the loop was the most flexible region of overall GFAT’s structure, as shown by root mean square fluctuation analysis (RMSF, Fig. 4A). During the simulation time, we observed that the loop approaches the protein itself in 2 replicas (Fig. 4B and 4C), but moves away from it in the 3^rd^ replica (Fig. 4D). By monitoring the network of interactions of loop residues, we noticed that Thr227, Asn230, Asn233, Arg238 and Arg241 are major players for the interaction with the interdomain region (mainly through residues Glu313 and Gln315, Fig. 5A-E and Suppl. Fig. 7A), and GLN (Glu269, Fig. 5A-D and 5F) and ISOM domains (Arg342 and Glu332, Fig. 5A-D, 5G and Suppl. Fig. 7B) in replicas 1 and 2.

**Fig. 4.**
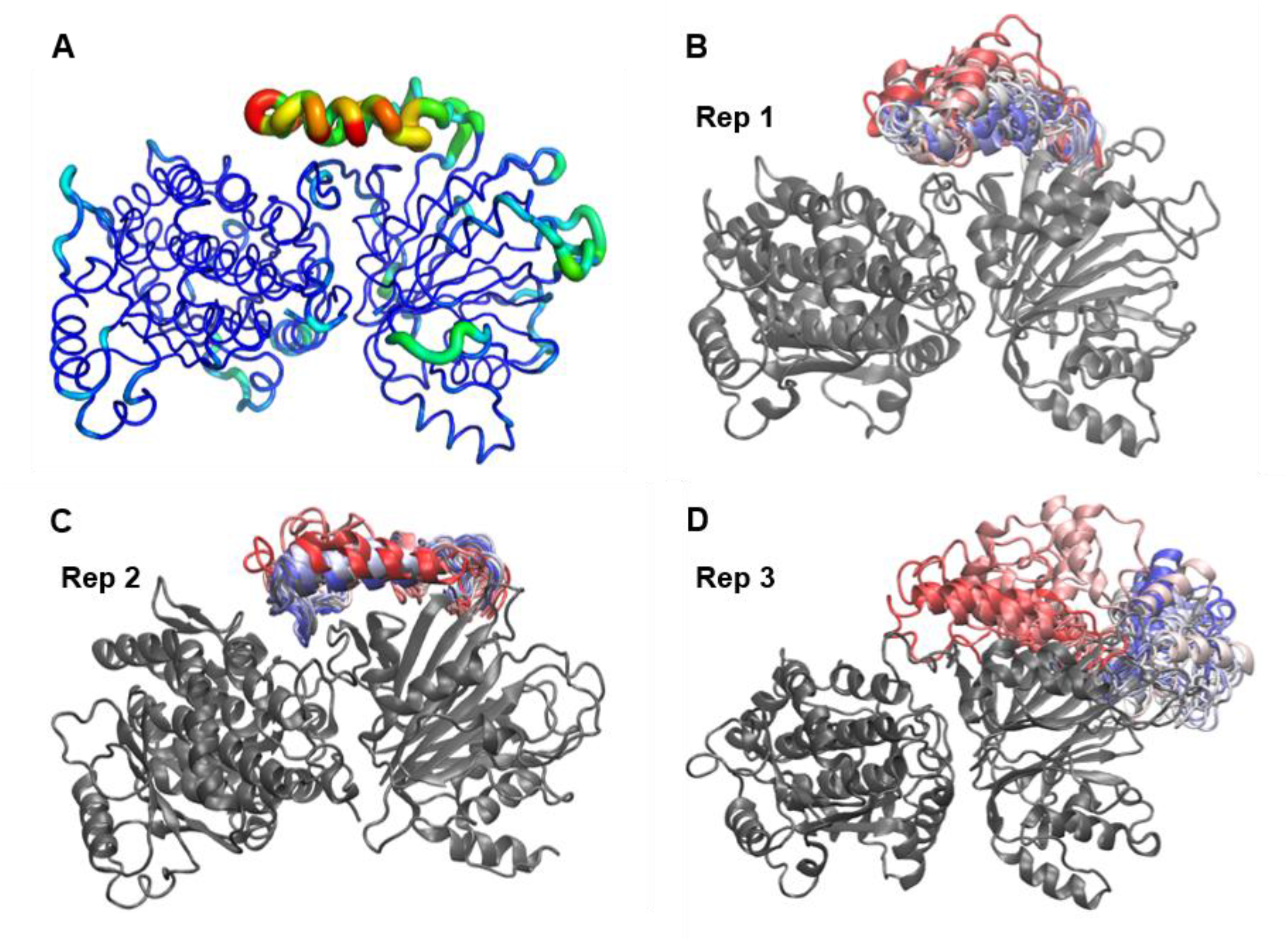
Loop stability. (A) Three-dimensional root mean square fluctuation (RMSF) of protein residues during MD simulation of replica 1. The protein is shown in tubes, whose thickness and color reflects the extent of each residue fluctuation (0.8 to 7.4 Å). (B-D) Final frames from MD simulation replicas, with the protein represented in cartoon and colored in gray. The loop conformation is shown throughout simulation time and colored accordingly: initial frames are colored in red, intermediates in white and final frames in blue.

**Fig. 5.**
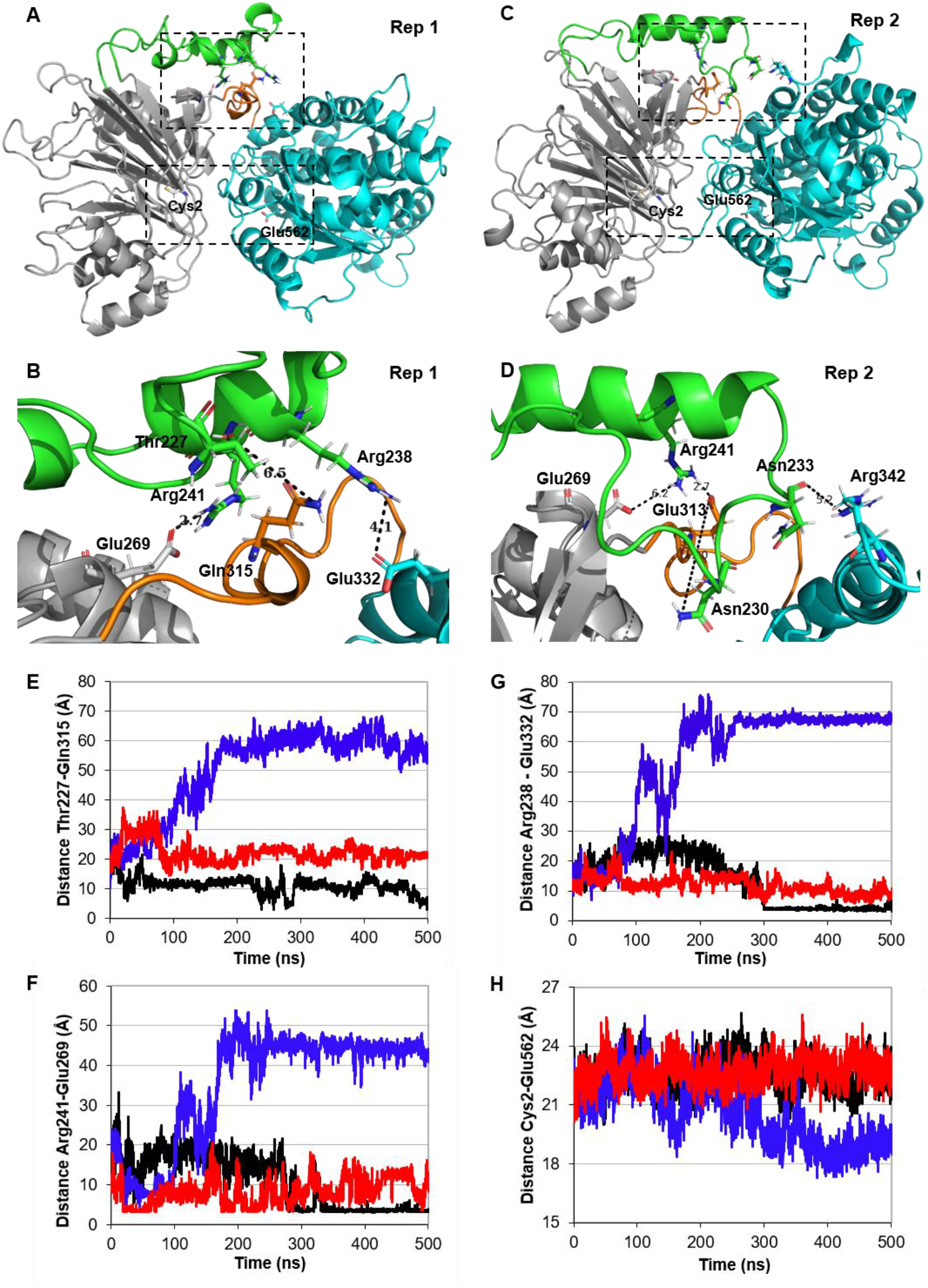
Structural features of rhGFAT2. (A-D) Views of rhGFAT2 from the most populated cluster from MD simulations of replicas 1 and 2, depicting the full protein structure (A and C) and closer views of the loop region (B and D). The protein structure is represented in cartoon and its regions are colored as follows: the GLN domain in gray, the loop in green, the ISOM domain in cyan and the interdomain region in orange. Key residues monitored in distance analysis are represented as sticks and the distances are represented by dashed black traces. (E-G) Analysis of the distances between the residues Thr227 and Gln315 (E), Glu269 and Arg241 (F), and Arg238 and Glu332 (G), reflecting the interaction of the loop with the interdomain region, the GLN domain, and ISOM domain, respectively, throughout simulation time. (H) Distance between catalytic residues from GLN domain (Cys2) and ISOM domain (Glu562) during MD simulation time. Black lines represent the data from replica 1, red lines from replica 2, and blue lines from replica 3.

Cluster analysis of MD frames from replicas 1 and 2 shows a heterogenous population distribution, in which few clusters – the ones reporting the loop in close contact with interdomain region and ISOM domain residues – accounts for more than half of the frames (Suppl. Fig. 7D), while the same analysis of replica 3 produced a greater number of less populated clusters (Suppl. Fig. 7D). These results indicate that the interaction between the loop and protein residues ensures its stabilization.

To assess whether the loop dynamics had effects on the movement of the domains, we monitored the distance between key residues from catalytic sites of both domains – Cys2, the suggested N-terminal nucleophile for glutamine hydrolysis on glutaminanse domain, and Lys559 and Glu562 (equivalent to Lys485 and Glu488 from GlmS) from ISOM domain – to assess whether the domains’ movement have an impact on them. In this way, we observed that, in the replicas in which the loop moves towards the protein (replicas 1 and 2), the domains did not move substantially, but in the replica in which the loop moves away from the protein (replica 3), they get closer by 4-5 Å (Fig. 5A-D, 5H and Suppl. Fig. 7C).

In order to understand how the structure of hGFAT2 could impact the ammonia leakage, we evaluate the neighborhood of Trp93 (unique Trp residue in this protein), equivalent to Trp74 in GlmS. Even though we observed conserved interactions between Trp93 and residues from Q-, R- and C-loops – such as Tyr35, Leu675, Ala676, and Arg33 (Fig. 6A-C) – the C-tail is oriented upwards relative to that observed in GlmS structure bound to DON and Glc-6P (Fig. 6D). In hGFAT2, this feature seems to be derived from the interaction between the loop an interdomain region, and this last with R-loop. In Fig. 6E we can observe that the hydrogen bonds between Arg29 and side chains and backbone from residues Glu313, Leu314 and Gln316 forces a turn in the interdomain connection. In contrast, Arg22 in GlmS, although close to Tyr240, seemed not to form a hydrogen bond to such residue, nor to any other within the interdomain region (Fig. 6F). Moreover, the interdomain region sequence is distinct and three residues longer in both hGFAT1 and hGFAT2 compared to GlmS (Fig. 3), which can extend its structure linearly up to 9.6 Å. Together, these are the first enzymatic and structural data from the hGFAT2.

**Fig. 6.**
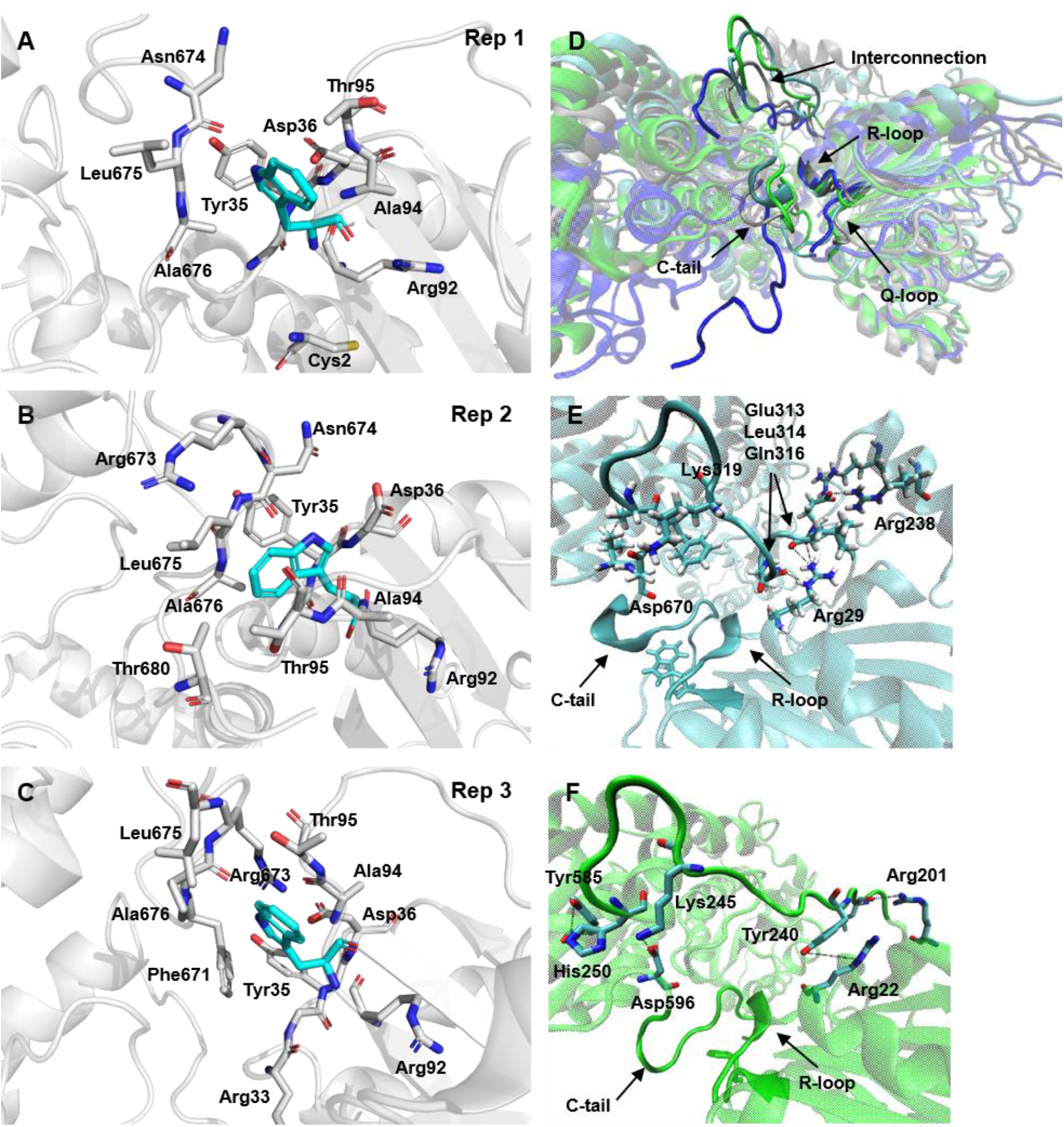
Tryptophan neighborhood in hGFAT2 and GlmS. (A-C) Closer view of the residues that are 4 angstroms from Trp93 on the most populated cluster from MD simulations of replicas 1 (A), 2 (B), and 3 (C). The residues in contact to Trp93 are represented in sticks and colored in gray; Trp93 is also in sticks but colored in cyan (the hydrogens were removed for clarity). The protein backbone is represented in cartoon and colored in gray. (D) Views of the most populated cluster from MD simulations of replicas 1 (gray), 2 (blue), 3 (cyan), aligned to GlmS structure (green, PDB ID: 2J6H). The proteins were aligned by GLN domain and are represented in cartoon; the interdomain region, C-tail and Q- and R-loops are highlighted and indicated. (E, F) Closer view of the interdomain region in replica 2 of hGFAT2 DM (E) and GlmS (F), depicting the interactions between this region and the overall protein. The residues involved in those interactions are represented in sticks and colored in cyan, and the distances are represented by dashed black traces.

## Discussion

Despite the involvement of hGFAT2 in cancer aggressiveness, few studies have focused on the molecular and structural characterization of this protein. Here, we conducted a comprehensive study detailing the enzymatic properties of hGFAT2. We first expressed and purified the rhGFAT2 either fused or not to a HisTag at its C-terminal end. We showed that this enzyme mostly forms tetramers, in agreement with the data from hGFAT1 His6-Asn298 (21), and also forms higher order oligomeric structures, possibly octamers, at lower extent. Even though the presence of higher order oligomeric structures was not reported so far for eukaryotic GFATs, the formation of octamers could exert some regulatory role for hGFAT2, since the interplay between quaternary structures has been described for *E. coli* GlmS as a mechanism of enzyme inactivation (30).

Regarding the enzymatic properties of hGFAT2, the fitting of rate curves from GlcN-6P synthetic activity using the Michaelis-Menten equation resulted in ^*app*^*K*_*M*_ for Fru-6P and Gln similar to those described for mGFAT2 (24), hGFAT1 with no tags (23), and hGFAT1 His6-Asn298 (21,22), but the and ^*app*^*K*_*cat*_ were lower than those reported for either mammalian and prokaryotic GFATs (20,21). When we fitted the data to ordered bisubstrate mechanistic model, based on kinetic studies with GlmS, we obtained an inconsistent value for ^*Fru-6P*^*K*_*M*_, suggesting that hGFAT2 does not follow such kinetic model. This perspective is corroborated by the aminohydrolysing data, which shows that Gln is hydrolyzed by GFAT even in absence of Fru-6P, whereas the addition of this monosaccharide-phosphate increases the glutaminase catalysis by 4-fold. Such a feature was also observed for GFA from *C. albicans* (31). Altogether, these data suggest that Fru-6P binding is not essential for Gln binding, but acts as an activator of GFAT2’s glutaminase activity. Since similar pattern of Fru-6P activation of aminohydrolysing activity was observed to rhGFAT2 without HisTag, we are confident that this phenomenon – the non-conditioning pattern of Fru-6P for Gln binding – is not an artifact derived from the HisTag. Although our data were not enough to determine the proper kinetic model that fits hGFAT2 GlcN-6P synthetic activity, it strongly indicates that this enzyme does not follow the ordered bi-bi substrate model, in contrast to the GlmS (19).

NMR spectroscopy was used to directly detect the progress of hGFAT2 catalysis. We found a considerable amount of Glc-6P, suggesting that hGFAT2 can partially act as an isomerase-only enzyme, regardless of Gln presence. This data is corroborated by the isomerase assays with rhGFAT2 either with or without HisTag. The phosphoglucose isomerase (PGI)-like activity has already been reported for GlmS (32) and for GFA in the absence of Gln (31). However, it is the first data describing that a human GFAT retains such a PGI-like activity, and moreover, that this activity is maintained even in the presence of Gln. Our findings are in agreement with previous report detecting Glc-6P upon cocrystalyzation of hGFAT1 with Gln and Fru-6P (22).

The isomerase data indicate, in the other hand, that part of Gln is lost in an unproductive hydrolysis. Indeed, the ammonia release assays suggest that Fru-6P does not prevent the loss of NH3 to the medium. To measure the efficiency of ammonia transfer, Floquet et al. (33) used the ratio between the *kcat* of synthase and hemi-synthase (glutaminase) activities, which, in their work, is 84% for GlmS. We and Ruegenberg et al. (22) observed an efficiency of ammonia transfer of approximately 50 and 47% for hGFAT2 and hGFAT1, respectively, indicating that human GFATs, in fact, have a higher rate of ammonia leakage. Structural data from GlmS point to the formation of a hydrophobic channel formed among Trp74 in the Q-loop and C-tail residues upon the binding of the two substrates as a key event for avoiding ammonia leakage (33–35). The movement of the Q-loop upon substrates binding was also observed for hGFAT1 (22), but it did not prevent the low efficiency of ammonia transfer. In this regard, kinetic and mutagenesis data from GFA suggest that deletion of a sequence from GLN domain disrupts the communication of both domains and hampers the GlcN-6P synthesis, but retains their aminohydrolysing and isomerase-only activities (31). In the present work, we observed that sequence from GFA is present in both hGFAT1 and hGFAT2, and folds in a highly flexible loop in the latter. The flexibility of this loop was also reported for hGFAT1 (22) and may be the reason for the difficulty in getting crystals from eukaryotic GFATs, as noticed by Nakaishi et al. (36).

Our MD simulation data suggest that the loop alternates among what could be seen as conformational states, in which it interacts with the interdomain region and ISOM domain, or shifts to an open-conformation. These results are in line with GlmS data from MD simulations and normal mode analysis (37). The hinge movement, which is reported to be performed solely by the hinge connection residues in GlmS, may be modulated by the loop residues in hGFAT2, and possibly in hGFAT1. In addition, the hinge connection sequence is distinct and is three residues longer in both hGFAT1 and hGFAT2 compared to GlmS, which could alter the domains’ motions and impact the sealing of hydrophobic channel by changing the orientation of R- and Q-loops to C-tail. Thus, our data suggest that the loop evolved as an additional regulatory mechanism, which is corroborated by having conserved phosphorylation sites in GFA (31,38), GFAT from *Drosophila melanogaster* (39), mGFAT2 (24) and hGFAT1 (40–42), a post-translational modification that alter their enzymatic properties with direct impact in cell biology (43).

The differences among hGFAT1 and hGFAT2 goes beyond the hinge connection and loop sequences, or their catalytic efficiency: they also differ in susceptibility to allosteric inhibition by UDP-GlcNAc (20,22). Our data is in accordance with previous work describing the partial inhibition of mGFAT2 by UDP-GlcNAc (24). The close interaction observed between the loop residues and Arg342, near the allosteric site, could lead to pocket hidrance and may explain the poor inhibition.

Despite these number of contrasting properties of hGFAT2 and hGFAT1, it is worth noting that both isoforms are simultaneously expressed in the vast majority of cell types so far analyzed, although their ratio vary among them (7,18,25). Moreover, several evidences have shown that their expression is modulated by different factors (Xpb1s for hGFAT1 and NR4A1 for hGFAT2, for example) (25,44) and is observed in distinct circumstances (18,45,46). This sheds light on the relevance of the difference between the characteristics of these two enzymes, suggesting that cells can take advantage of it by changing the ratio hGFAT1/hGFAT2. In addition, we cannot exclude the possibility that hGFAT1 and hGFAT2 form heterotetramers. Altogether, our work provides the first comprehensive set of data on the structure, kinetics and mechanics of hGFAT2. Our results contribute to the knowledge of physiological roles and differences between GFAT isoforms. More studies addressing the interaction of hGFAT2 to substrates and ligands are important.

## Experimental procedures

### Construction of pET-hGFAT2 plasmids

The *gfpt2* gene was amplified from pCMV6-AC plasmid (Origene, USA) by PCR and inserted into bacterial expression plasmid pET23a (Novagen, USA) in order to construct either pET-hGFAT2 with and without HisTag. For both plasmids, the gene amplification was performed using the same sense primer 5’ GGAATTCCATATGTGCGGAATCTTTGCCT AC 3’ (containing a restriction site for Nde I), but distinct antisense primers: 5’ ATAAGAATGCGGCCGCTTCCACAGTTACAGACTTG 3’ for hGFAT2-His, and 5’ ATAAGAATGCGGCCGCTTATTCCACAGTTACAGACTTG 3’ for hGFAT2 without tag (both containing a restriction site for Not I), this last containing a stop codon right after the protein sequence. The reactions were performed as follows: 2 min at 94 °C followed by 35 cycles of 1 min at 94 °C, 1 min at 52 °C, and 2 min at 68 °C, with an extension step of 7 min at 68 °C. The amplified genes were then electrophoresed in 1% agarose gel followed by purification using PCR purification kit (Qiagen, USA). Both purified genes and plasmids were digested with Nde I and Not I prior to ligation using the T4 DNA ligase (New England Biolabs, UK). The recombinant plasmids pET-hGFAT2 (without tag) and pET-hGFAT2-his were inserted into electro-competent *E. coli* DH5α cells and positive colonies were subjected to a PCR colony using the abovementioned primers. The reactions was performed as follows: 2 min at 94 °C followed by 35 cycles of 1 min at 94 °C, 1 min at 52 °C, and 2 min at 72 °C, with an extension step of 7 min at 72 °C. True positive clones were isolated and sequenced by using an ABI PRISM dye terminator cycle sequencing core kit (Applied Biosystems, USA).

### Expression of recombinant hGFAT2s (rhGFAT2s) in *E. coli*

Chemically competent *E. coli* Codon plus cells (Novagen, USA) were transformed with 200 ng of the pET-hGFAT2 (with or without tag) plasmids, and positive clones were selected in an LB-agar medium containing 100 μg/mL ampicillin and 34 μg/mL chloramphenicol at 37 °C overnight. A single positive colony was pre-inoculated in 10 mL of LB medium containing 100 μg/mL ampicillin and 34 μg/mL chloramphenicol, and this culture was stirred at 220 rpm at 37 °C overnight. The overnight culture was diluted to 1:50 in 1 L of fresh antibiotic-containing medium and grown at 37 °C until an optical density (O.D._600nm_) of approximately 0.7-0.8 was reached. The induction of protein expression was conducted with 0.5 mM IPTG followed by 6 h of expression at 25 °C with 220 rpm stirring. Thus, the cells were harvested by centrifugation at 5,000 × *g* for 20 min at 4 °C, and the total-cell lysate was prepared.

### Purification of rhGFAT2s

The pellet was suspended in 25 mL of Buffer A (20 mM Tris-HCl pH 7.5, 500 mM NaCl, 1 mM DTT and 0.5% NP-40) in the presence of 1 mM PMSF and 0.5 μg/mL of each protease inhibitor: aprotinin, bestatin, pepstatin and E-64 (Sigma Aldrich, USA). Then, 5 mg/mL of lysozyme, 10 μg/mL of DNase A, and 5 mM of magnesium chloride were added, and the solution was incubated for 30 min at 4 °C with stirring. The total-cell lysate was sonicated using 10 cycles of 15 sec on and 1 min off at 40% amplitude, followed by centrifugation at 37,200 × *g* for 20 min at 4 °C.

The supernatant fraction containing the rhGFAT2 protein (with or without tag) was subjected to purification using a Ni^+2^NTA affinity column (HisTrap HP 5 mL, GE Healthcare, USA). The column was equilibrated with 10 column volumes (CV) of Buffer A prior to load the sample at a flow of 1 mL/min. After this step, the nonspecific ligands were removed by washing the column with 5 CV of Buffer A. The elution was performed using a gradient of Buffer A and Buffer B (Buffer A with the addition of 0.5 M imidazole) at a flow rate of 2 mL/min. All collected samples were analyzed by SDS-PAGE, and the tubes containing the purified rhGFAT2-his were pooled and dialyzed against Storage buffer (20 mM Tris-HCl pH 7.5, 150 mM NaCl, 1 mM DTT, and 5% glycerol).

The purification of rhGFAT2 w/o tag required an additional step of purification with an anion exchange chromatography. The SDS-PAGE analyzed fractions from HisTrap column which contained the rhGFAT2 w/o tag were pooled and dialyzed overnight against buffer C (20 mM Tris-HCl pH 8.0, 1 mM DTT, 0.5% NP40) with 150 mM NaCl. The dialyzed protein was diluted in buffer C to reach 50 mM NaCl immediately before loading to a Q-sepharose HP column (5 mL, GE Healthcare, USA), previously equilibrated with 10 CV of Buffer C. The nonspecific ligands were removed by washing the column with 5 CV of Buffer C. The elution was performed using a gradient of Buffer C and Buffer D (Buffer C with the addition of 0.5 M NaCl) at a flow rate of 2 mL/min. All collected samples were analyzed by SDS-PAGE, and the tubes containing the purified rhGFAT2 w/o tag were pooled and dialyzed against Storage buffer.

### Western Blot

The purified proteins (2 ug each) were submitted to SDS-PAGE in 10% acrylamide gel, and electrofransfered to nitrocellulose membrane. The membrane was blocked with 3% (w/v) bovine serum albumin in Tris-buffered saline with 0.1% (v/v) Tween-20, and incubated overnight at 4 °C with anti-His (Santa Cruz Biotechnologies, USA). The membrane was then washed, incubated for 1 h under agitation with the secondary antibody (anti-mouse, Santa Cruz). After a second round of washing, the labeled membrane was developed with Femto ECL (Thermo Fisher Scientific) and exposed to ImageQuant LAS 500 (GE Healthcare). The membrane was stripped and labelled with anti-GFAT2 (Cell Signaling Technologies, USA) following the same procedure described above.

### Cross-linking assay

The purified rhGFAT2 protein (3, 5 and 10 μg) was incubated in PBS buffer in the presence or absence of 1 mM EGS for 30 min at room temperature. The reactions were stopped with addition of 30 mM of Tris-HCl pH 8.0. Approximately 10 μg of each sample were analyzed by a gradient SDS-PAGE assay (Bio-Rad, USA) followed by Coomassie Brilliant Blue staining.

### Size exclusion chromatography

The purified rhGFAT2-his protein was subjected to a size exclusion chromatography using a Superdex 200 column (GE Healthcare, USA). The column was equilibrated with 1 CV of 20 mM Tris-HCl pH 7.5, 150 mM NaCl, 1 mM DTT, 0.5% NP-40 prior to sample loading at a flow rate of 1 mL/min. The fractions were collected and analyzed by SDS-PAGE. The molecular weight of rhGFAT2 oligomer was estimated according to the retention time of standard proteins (Thyroglobulin - 669 kDa, Apoferritin - 443 kDa, β-amilase - 200 kDa, BSA - 66 kDa, Carbonic anhydrase - 29 kDa, and Citocrome C oxidase - 12.4 kDa) acquired from Sigma Co.

### Characterization of rhGFAT2-his products by NMR

Solution of rhGFAT2-his was exchanged with deuterated sodium phosphate buffer (50 mM pH 7.4, with 150 mM NaCl and 1 mM DTT) using four cycles of dilution and concentration with Amicon Ultra 30K NMWL (Millipore, USA). Two hundred microliters of 100 μg protein solution were incubated with 3 mM of Fru-6P and 3 mM of Gln in Shigemi tubes. In order to check for spontaneous product formation or substrate consumption, the same amounts of Gln and Fru-6P were incubated with 200 μL of the deuterated buffer in which the protein were conditioned. NMR spectra were obtained at a probe temperature of 298 K on a Bruker Avance III 500 MHz equipped with a 5 mm self-shielded gradient triple resonance probe. The GFAT reaction products were monitored by unidimensional ^1^H spectra, performed according to the Bruker pulse sequence zgesgp. The product characterization was assisted by total correlation spectroscopy (TOCSY) spectra, which were recorded using mlevesgpph pulse sequence with a mixing time of 80 ms and 64 scans per t1 increment. For each scan, 8192 transients of 256 complex data points were acquired to a 10.0 ppm spectral width. The spectra were multiplied with a square cosine bell function in both dimensions and zero-filled two-fold. The data acquisition and analysis were performed using spectrometer software Topspin 3.6 (Bruker Corporation).

### Enzyme assays

#### GlcN-6P Synthetic activity

The specific GlcN-6P synthetic activity from rhGFAT2-his and rhGFAT2 w/o tag was assayed by incubating 100 μg of each protein with 10 mM Fru-6P, 10 mM Gln, 1 mM DTT in PBS pH 7.4 (100 μL of final reaction volume) for 1 h at 37 °C under agitation. The glucosamine-6-phosphate (GlcN-6P) formed in the reaction mixtures was determined as described by Queiroz et al. (27), based on Elson & Morgan (26). Briefly, 10 μL of 1.5% acetic anhydride (Sigma, USA) and 50 μL of 100 mM sodium tetraborate was added to the reaction mixture and incubated at room temperature for 5 min under agitation. The samples were then incubated at 80 °C for 25 min, cooled down at 4 °C for 5 min, and spun down for removing precipitated protein. The resultant acetylated GlcN-6P was derivatized with 130 μL of Ehrlich reagent in a 96-wells microplate incubated for 30 min at 37 °C and finally read at 585 nm in microplate reader (SpectraMax 190, Molecular Probes, USA). The absorbance of the samples not incubated with GFAT substrates was discounted and the concentration of GlcN-6P was determined comparing the resultant absorbance of the samples with GlcN-6P standards processed in the same manner. The specific activity was expressed as units (μmol of GlcN-6P synthesized per min at 37 °C) per mg of protein.

For kinetic analysis, the assay was performed as described, but with the following modifications: rhGFAT2-his was incubated with variable concentrations of one of the substrates (Fru-6P or Gln, at 0.156, 0.313, 0.625, 1.25, 2.0, and 2.5) while the other was fixed at saturating concentration (10 mM); the reaction mixtures were incubated by multiple time points up to 15 min, counted from the addition of the enzyme (time point 0 min was considered as the reaction mixture without the enzyme). At the end of incubation time, the reaction mixtures were processed for GlcN-6P derivatization as for the specific activity. For calculation of kinetic parameters, the progression curves were plotted, and the initial velocity was calculated (related to each substrate). The apparent kinetic parameters (*k*_*cat*_ and *K*_*M*_) were determined by direct fit of the rate *versus* substrate concentration data to the rate equation for Michaelis-Menten using GraphPad Prism version 8 (GraphPad, USA). The data were also submitted to fitting to both simple Michealis-Menten equation and ordered bisubstrate mechanistic equation using GraFit version 7 (Erithacus Software, USA).

#### Aminohydrolysing (glutaminase) activity

GFAT glutaminase activity was determined using a coupled assay, based on Ye et al.(47). In the assay, the glutamate released by GFAT activity is oxidized by glutamic acid dehydrogenase (GDH) with concomitant 3-acetylpyridine adenine dinucleotide (APAD) reduction. The amidotransferase reaction was carried out in 200 μL of 20 mM phosphate buffer pH 7.4 with 50 μg of rhGFAT2-his, containing variable concentrations of Gln (0.156, 0.313, 0.625, 1.25, 2.5, 5.0, and 10.0 mM), and Fru-6P. APADH formation was monitored continuously by absorbance at 370 nm in Spectramax 190 instrument (Molecular Devices, CA, USA) for 1 h at 37 °C. The APADH concentration was derived from its molar extinction coefficient. Kinetic parameters were determined as described for GlcN-6P synthetic activity.

#### Isomerase activity

The isomerization of Fru-6P to Glu-6P by rhGFAT2 was assayed as described by Olchowy et al. (31). In brief, 50 μg of rhGFAT2-his were incubated with variable concentrations of Fru-6P in 200 μL of 50 mM Tris-HCl pH 7.4 with 1 mM DTT, 0.5 mM NADP (Sigma, USA) and 2.5 mU/μL glucose-6-phosphate dehydrogenase (G6PD from *Saccharomyces cerevisiae*, Sigma, USA). Some assays were performed in the presence of variable concentrations of Gln (0.5, 0.625, 1.25, 2.5, 5.0 and 10 mM) with fixed (0.625, 2.5 and 10 mM) concentrations of Fru-6P. NADPH formation was monitored continuously by absorbance at 340 nm in Spectramax 190 instrument (Molecular Devices, CA, USA) for 30 min at 25 °C. The NADPH concentration was derived from its molar extinction coefficient. Kinetic parameters were determined as described for GlcN-6P synthetic activity.

To evaluate the ISOM specific activity of rhGFAT2 with or without HisTag, each enzyme was incubated with 10 mM Fru-6P in the presence or absence of 10 mM Gln in the same conditions described above. The absorbance at 340 nm was read at the end of 30 min. The specific ISOM activity was expressed as units (μmol of NADPH synthesized per min at 37 °C) per mg of the enzyme.

#### Ammonia release

The release of ammonia from Gln hydrolysis catalyzed by rhGFAT2 was monitored by using the GDH in reverse direction, based on Floquet et al. (33). In this perspective, the ammonia released by GFAT activity is used in reductive amination of α-ketoglutaric acid (αKG), thereby with NADH oxidation. The assays were carried out by incubating 50 μg of rhGFAT2 (with or without HisTag) with variable concentrations of Gln (0.25, 1.25 and 10.0 mM), or with a fixed saturating concentration of Gln (10 mM) and variable concentrations of Fru-6P (0.25, 1.25, 2.5, 10, and 20 mM), in 200 μL of 20 mM phosphate buffer pH 7.4 with 1 mM DTT, 0.25 mM NADH, 2.5 mM αKG, and 30 mU/μL GDH. Reaction mixtures with the enzymes and the variable concentrations of Gln (0.25, 1.25 and 10.0 mM), but without αKG were used as blanks for their correspondent reactions. The NADH consumption was assessed by absorbance at 340 nm in Spectramax 190 instrument after 30 min incubation at 37 °C. The NADH consumed was taken as the difference between the final absorbance of each of the samples and their correspondent blanks without αKG. The ammonia released was determined comparing the resultant absorbance difference of the samples with the ones from NH_4_Cl standards. The results were expressed as units μmol of ammonia per min per mg of protein.

### Peptide fingerprinting

Five micrograms of rhGFAT2-his was reduced with 3 mM DTT at 60 °C for 30 min, and carbamidomethylated with 9 mM iodoacetamide at room temperature for 30 min in the dark. The protein was then digested with Trypsin Gold (Promega) 1:100 at 37 °C overnight in 10 mM ammonium bicarbonate pH 8.0 and the resultant peptides were cleaned up with POROS 20 R2 (Applied Biosystens). The sample was dried under vacuum, solubilized in 2% acetonitrile and 0.1% formic acid (FA) in water and submitted to LC-MS in Nexera UPLC system (Nexera, Shimadzu, Japan) coupled to maXis Impact mass spectrometer (Q-TOF configuration, Bruker Daltonics) equipped with electrospray ionization source. Separation was accomplished in an Acquity CSH C18 UPLC column (150 m × 1 mm, 1.7 μm particle size, Waters) at 50 °C using a flow rate of 130 μL/min. After equilibration with 0.1% formic acid in water containing 2% acetonitrile, the peptides were injected and eluted using the following acetonitrile gradient: 2-8% in 2 min, 8-25% in 28 min, 25-50% in 10 min and kept at 50% for 2 min, 50-95% in 1.5 min and kept at 95% for 6 min. The electrospray source parameters were set as following: capillary voltage at 4.5 kV, end plate offset at −500 V, nebulizer gas at 1.2 bar, dry gas at 8 L/min, and dry temperature at 200 °C. Mass spectra were acquired in the positive-ion mode over the range *m/z* 50–1500 in data dependent acquisition fragmentation mode at 1 Hz. The mass spectrometer was internally calibrated using 100 μM sodium formate solution.

The mass spectrometry data was processed using Mascot Search engine (Matrix Science) in BioTools software version 3.2 (Bruker Daltonics). The MS/MS data were searched against both the Uniprot Human amino acid sequence database and the hGFAT2 sequence, with and without Met1, for protein/peptide identification. The search was set up for full tryptic peptides with a maximum of 2 missed cleavage sites; carbamidomethyl cysteine and oxidized methionine were included as fixed and variable modifications, respectively. The precursor mass tolerance was set to 10 ppm, and the maximum fragment mass error was set to 0.05 Da.

### Circular Dichroism

The circular dichroism (CD) experiments were conducted with hGFAT2-his in a Chirascan Circular Dichroism Spectropolarimeter (Applied Photophysics, UK) at 20 °C using a quartz cuvette with a 0.01 cm path length. Spectra from three scans from 260 to 190 nm at a 30 nm/min speed were averaged, and the buffer baselines were subtracted from their respective sample spectra. As a negative control, the protein was further denatured with 6 M guanidine-HCl and CD scans were repeated. The secondary structure content was estimated from fitting the far-UV CD spectra using the different algorithms, such as CDSSTR, K2D (48) and SELCON3 (49,50), which is available on the Dichroweb server (51,52).

### Modelling GFAT structures

Multiple sequence alignments of hGFAT2 and its orthologous enzymes were carried out using ClustalO (53). The hGFAT2 sequence was submitted to the I-TASSER (Iterative Threading Assembly Refinement) (54) server to achieve a complete structural model. The hGFAT1 and GFA (from *C. albicans*) sequences were submitted to I-TASSER server as well. The best models were selected based on the higher confidence scores and template modeling scores.

### Molecular dynamics simulation

The best hGFAT2 model was further submitted to molecular dynamics simulation to investigate its conformational stability. Molecular Dynamics (MD) simulations were performed using the AMBER v. 14 software package (55) with the AMBER ff14SB force field (56). Explicit TIP3P water molecules (57) were used to solvate the hGFAT2 structure model in a cubic water box, using periodic boundary conditions. The protonation state of protein residues was assigned according to the values at pH 7.4 using the PROPKA software (58). The system was then neutralized by adding 1 Na^+^ ion to the simulation box. SHAKE algorithm (59) was applied to constrain all the bonds involving hydrogen atoms. Long-range electrostatic interactions were calculated with the PME method (60). The nonbonded interactions (Coulomb and van der Walls) were calculated using cutoffs of 8 Å.

The system was energy-minimized using 25000 cycles of Steepest Descent algorithm followed by 25000 cycles of Conjugated Gradient method with and without position restraint of 5 kcal mol^−1^ Å^−2^ for protein heavy atoms. The system was gradually heated from 0.15 to 300 K over 200 ps. Langevin thermostat (61) with a collision frequency of 0.067 ps^−1^ was used to control the temperature under a canonical ensemble and applying positional restrictions to the protein heavy atoms. Next, the pressure was applied until stabilized at 1 bar, using Berendsen barostat (62), by 7.5 ns under an isothermal and isobaric MD simulation with protein heavy atoms restrained to adjust the solvent density. The force constant for restraint was decreased gradually from 3 to 0 kcal^−1^ Å^−2^. Finally, 500 ns of production MD simulation with a time step of 2 fs was performed at a constant temperature of 300 K using Langevin themostat with a collision frequency of 5.0 ps^−1^ and a constant pressure of 1 bar controlled by Berendsen barostat (62) with a 1 ps pressure relaxation time. The MD trajectory was saved every 100 ps for analysis. The MD simulations replicas were performed by using different seeds to generate initial velocities. The analysis and figures were made using PyMol (The PyMOL Molecular Graphics System, Version 1.2r3pre, Schrödinger, LLC.) and VMD (63) programs.

## Abbreviations

αKG: α-ketoglutaric acid;
APAD: 3-acetylpyridine adenine dinucleotide;
APADH: reduced form of APAD;
EGS: ethylene glycol bis(succinimidyl succinate);
Fru-6P: fructose-6-phosphate;
G6PD: glucose-6-phosphate dehydrogenase;
GDH: glutamic acid dehydrogenase;
GFAT: glutamine:fructose-6-phosphate amidotransferase;
GFA: glutamine:fructose-6-phosphate amidotransferase from C. albicans;
Glc-6P: glucose-6-phosphate;
GlcN-6P: glucosamine-6-phosphate;
GlmS: glucosamine-6-phosphate synthase from E. coli;
GLN: glutaminase (in reference to protein domain or activity);
HBP: hexosamine biosynthetic pathway;
hGFAT: human GFAT;
IPTG: isopropyl-β-D-thiogalactoside;
ISOM: isomerase (in reference to protein domain or activity;
LC-MS: liquid chromatography coupled to mass spectrometry;
MD: molecular dynamics;
mGFAT: murine GFAT;
PGI: phosphoglucose isomerase;
rhGFAT: recombinant human GFAT;
RMSF: root mean square fluctuation;
TOCSY: total correlation spectroscopy.

## Acknowledgements

We thank Centro Nacional de Ressonância Magnética Nuclear (CNRMN - CENABIO, UFRJ), Centro de Espectrometria de Massas de Biomoléculas (CEMBIO, UFRJ), Plataforma de Expressão e Purificação de Proteínas com Interesse Biotecnológico (PEPIBiotec, UFRJ), and Plataforma de Imunoanálise (PIA, UFRJ). T.V.A.F. acknowledges Instituto Nacional de Metrologia, Qualidade e Tecnologia (Programa Nacional de Apoio ao Desenvolvimento da Metrologia, Qualidade e Tecnologia - PRONAMETRO) for a scholarship.

